# BULLKpy: An AnnData-Inspired Unified Framework for Comprehensive Bulk OMICs Analysis

**DOI:** 10.64898/2026.01.26.701768

**Authors:** Marcos Malumbres

**Author notes:** Correspondence:* M. Malumbres, Vall d’Hebron Institute of Oncology (VHIO), Natzaret 115-117-CELLEX, 08035 Barcelona.

## Abstract

Bulk OMICs data, such as RNA-seq and proteomics, remain foundational in biomedical and cancer research. While single-cell transcriptomics has revolutionized our understanding of cellular heterogeneity, bulk OMICs continue to provide a solid ground for large-scale studies, integrative analyses, and robust biomedical correlations. Yet, analytical workflows for bulk OMICs are frequently fragmented across disparate tools, programming languages, and data formats, in contrast to the single-cell field, which benefits from comprehensive and standardized frameworks like Scanpy and the broader scverse ecosystem. Here, we introduce BULLKpy, a Python-based, scverse-inspired framework designed to deliver similar integration, flexibility, and scalability to bulk RNA-seq analysis, with a particular emphasis on cancer research. Beyond standard preprocessing and visualization utilities, BULLKpy offers systematic evaluation of gene-metadata associations, categorical enrichment analyses, clustering stability metrics, and advanced visualization strategies that facilitate intuitive exploration of tumor heterogeneity in expression profiles and metaprograms. Cancer-specific visualizations, such as oncoprints, are natively supported and seamlessly integrated with transcriptomic and clinical features, allowing users to relate somatic alterations to expression-derived phenotypes. By standardizing workflows within the Python ecosystem and aligning with AnnData objects and the scverse, BULLKpy aims to democratize advanced transcriptomic and proteomic analyses, facilitate integrative cancer research, and support future developments at the intersection of computational biology, multi-omics integration, and artificial intelligence.

## Introduction

High-throughput OMICs such as transcriptomics or proteomics have become foundational in biomedical research, spanning molecular classification, biomarker discovery, and clinical association studies. While single-cell technologies have transformed our ability to resolve cellular heterogeneity (Heumos *et al*, 2023; Stuart *et al*, 2019), bulk RNA sequencing (RNA-seq) or proteomics remains indispensable for large clinically annotated cohorts and retrospective outcome analyses, where sample sizes, standardized processing, and clinical metadata are often richer than in single-cell collections. In the Python environment, Scanpy and the broader scverse ecosystem (Virshup *et al*, 2023; Wolf *et al*, 2018) have played a defining role in single-cell analysis by establishing a coherent workflow built around a unified annotated data structure and modular tool namespaces for preprocessing, analysis, and visualization. This design has enabled scalable exploration, consistent reporting, and rapid method development, accelerating adoption across the community (Wolf *et al*., 2018). These frameworks have not only lowered the barrier to entry for complex analyses but have also promoted reproducibility and interoperability across studies and modalities. By contrast, bulk RNA-seq or proteomics analysis in Python is still commonly performed through heterogeneous, manually assembled pipelines that combine multiple libraries for normalization, differential expression, enrichment, and clinical modeling. Even when upstream processing is standardized by community workflows (e.g., nf-core/rnaseq), downstream cohort analysis often remains fragmented across scripts and toolchains, increasing variability and slowing iteration, which imposes a burden for a large number of scientists with more limited computing skills.

Cancer-focused OMICs studies also present recurring analytic requirements that are not consistently supported as first-class features in general-purpose toolkits. These include systematic gene–metadata association analyses (linking expression or signatures to clinical variables, molecular subtypes, or continuous phenotypes), deeper interpretation of enrichment results via identification of core driver genes shared across enriched pathways, and visual summaries of the correlation between expression profiles and genomic alterations in clinically stratified cohorts. Leading-edge subsets are central to interpreting gene set enrichment and pathway studies because they capture the genes most responsible for the enrichment signal and enable functional clustering and biological predictions.

To address these needs, we developed BULLKpy, a comprehensive Python framework for bulk OMICs analysis that adopts a Scanpy-inspired philosophy; i.e. a unified AnnData object coupled with modular, interoperable analysis and visualization functions. BULLKpy is designed for cohort-scale cancer transcriptomics or proteomics, integrating preprocessing and quality control, dimensionality reduction and structure exploration, gene signature scoring, differential expression, enrichment interpretation (including leading-edge overlap analyses), gene–metadata associations and other statistical studies, as well as cancer-centric visualization such as oncoprints. By consolidating common bulk OMICs tasks into a coherent API and emphasizing reproducibility and clinical relevance, BULLKpy aims to complement existing upstream processing pipelines and specialized libraries, while enabling state-of-the-art, standardized downstream analyses for translational oncology without the need of computing skills.

## Results

### Design principles and compatibility with the AnnData ecosystem

At its core, BULLKpy adopts the AnnData object (Virshup *et al*., 2023; Wolf *et al*., 2018) as its central data structure, enabling the joint storage of expression matrices, feature annotations, sample-level metadata, dimensionality reductions, graph representations, and analysis results (Figure 1). AnnData stores a data matrix (X) linked to annotations of observations (samples and their metadata) and variables (genes, proteins, features), and unstructured annotations. Major difference with the use of AnnData in single-cell studies is that samples represent tumors (or any other biological material) rather than single cells, and this determines specific algorithms and, mostly, different interpretations in some downstream analyses. Yet, this design choice ensures not only direct conceptual and technical compatibility with the scverse (Virshup *et al*., 2023), but also the seamless implementation of new modules and connection to existing pipelines (e.g. GSEApy; see below), and future connections to new algorithms that can eventually be applied to bulk data. Mirroring the Scanpy API (Wolf *et al*., 2018), BULLKpy organizes functionality into three primary modules: “pp” (preprocessing), “tl” (tools), and “pl” (plotting). The “pp” module implements standardized normalization, filtering, and quality-control routines tailored to bulk OMICs data. The “tl” module provides higher-level analytical methods, including dimensionality reduction, clustering, functional scoring, differential expression, gene set enrichment analysis, metaprogram inference, and survival modeling. The “pl” module generates publication-ready visualizations that are tightly coupled to the underlying AnnData object, ensuring consistency between analyses and figures (Figure 1). Structured entry data and metadata, as well as many of the results from BULLKpy “pp” and “tl” pipelines and internally stored in python within the AnnData object and can be save externally into.h5ad or similar high-structure files for sharing or future use.

**Figure 1.**
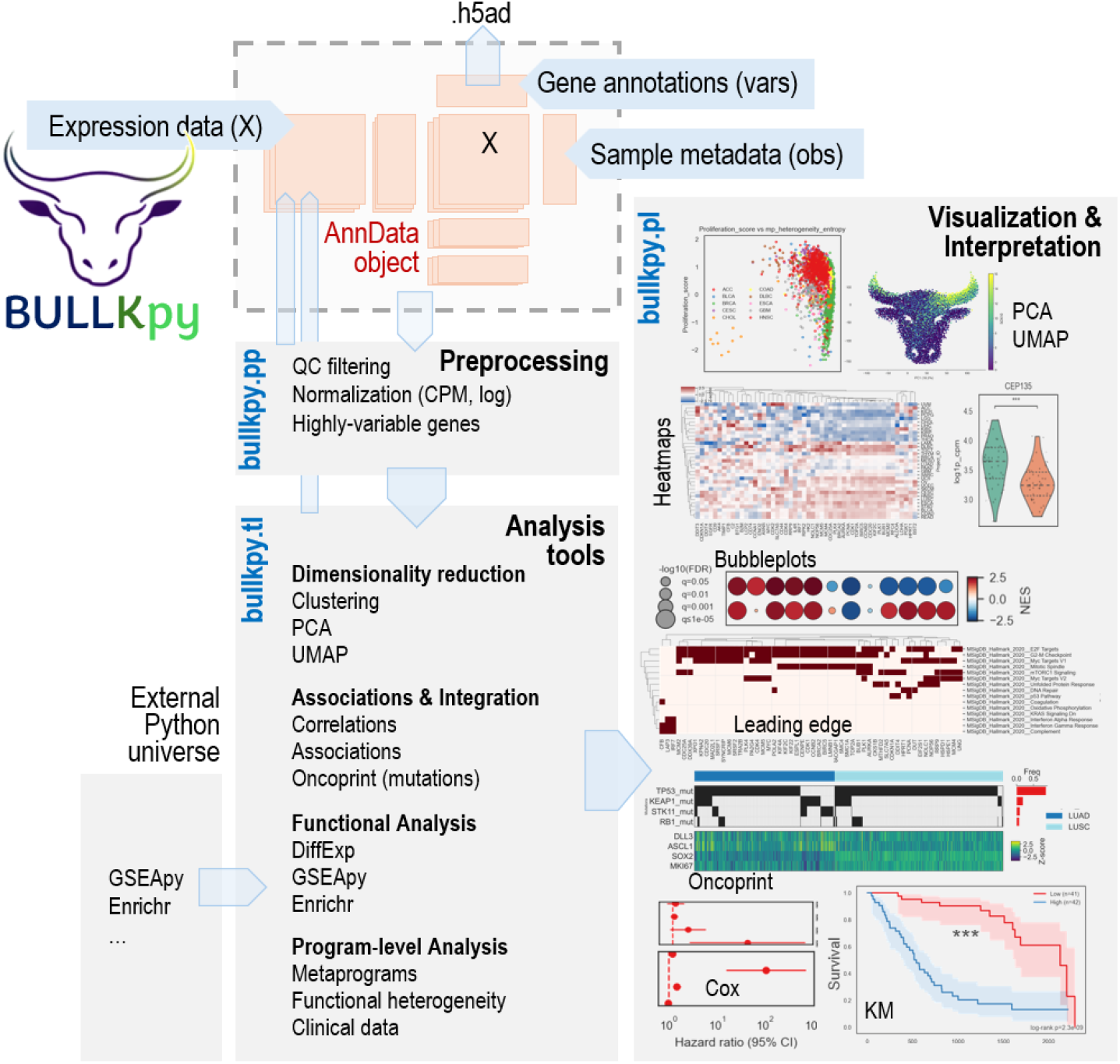
Overview of the BULLKpy analytical workflow and API. BULLKpy [import bullkpy as bk] adopts the AnnData data model and a Scanpy-like modular design, organizing bulk RNA-seq analyses into preprocessing (bk.pp), analytical tools (bk.tl), and visualization (bk.pl) modules. Raw expression matrices and sample or gene/protein metadata are imported into a unified AnnData dataset object, which is used throughout the analysis. Pre-processing includes quality control, filtering, normalization, and batch correction. Dimensionality reduction and structure inference enable exploration of sample relationships. Biological interpretation is supported through gene signature scoring and association analyses. Statistical and clinical modules facilitate differential expression, pathway enrichment, and survival modeling. This structure enables integrated analyses of bulk transcriptomic data with functional interpretation and clinical association, with a particular focus on cancer biology

By definition, bulk OMICs data are limited by the heterogeneity of the sample, including the presence of malignant and non-malignant (immune, stroma, etc.) in the case of cancer studies. Yet, most clinical studies typically lack single-cell data, and they are still largely relying in bulk analyses. In addition, bulk data can be easily combined with additional biological features, such as DNA mutations or clinical response to treatments, etc. that are typically not available for single cells or exist at the tumor or patient level. We therefore added to BULLKpy several modules that are linked to mutational studies or correlation with metadata or clinical data typically available at the sample/patient level.

### Preprocessing and bidimensional representation of data

To demonstrate the utility of BULLKpy and implement classical and new functions into an integrated framework, we made use of bulk RNA-seq data from the TCGA initiative (Cancer Genome Atlas Research *et al*, 2013) (Hoadley *et al*, 2018). Expression matrices and related metadata were obtained from public repositories (https://portal.gdc.cancer.gov/; https://xenabrowser.net/) or specific publications reporting further analysis of these data for specific features such as immune infiltration, aneuploidy or detailed clinical metadata (Liu *et al*, 2018; Saltz *et al*, 2018; Taylor *et al*, 2018).

We used in this work a matrix of 52,603 transcripts x 10,534 samples, representing 33 tumor types following the TCGA study abbreviations https://gdc.cancer.gov/resources-tcga-users/tcga-code-tables/tcga-study-abbreviations). These data were imported into the AnnData object and several quality control (QC) utilities were implemented following pipelines already available in scanpy for single-cell studies (Wolf *et al*., 2018), such as quantification of counts and genes detected, as well as scoring of mitochondrial genes, and pairwise QC visualizations as a readout of possible poor quality of the samples due to excessive cell death (Figure 2a,b). BULLKpy offers several possibilities and plots for QC not only in the complete dataset but also after grouping by specific categorical metadata (e.g. tumor type). After filtering out poor-quality samples or genes, normalization (cpm) and log-transformation (log1cpm) is next performed and these different matrices (counts, cpm and log1cpm) can be conserved in the AnnData object for possible later studies.

**Figure 2.**
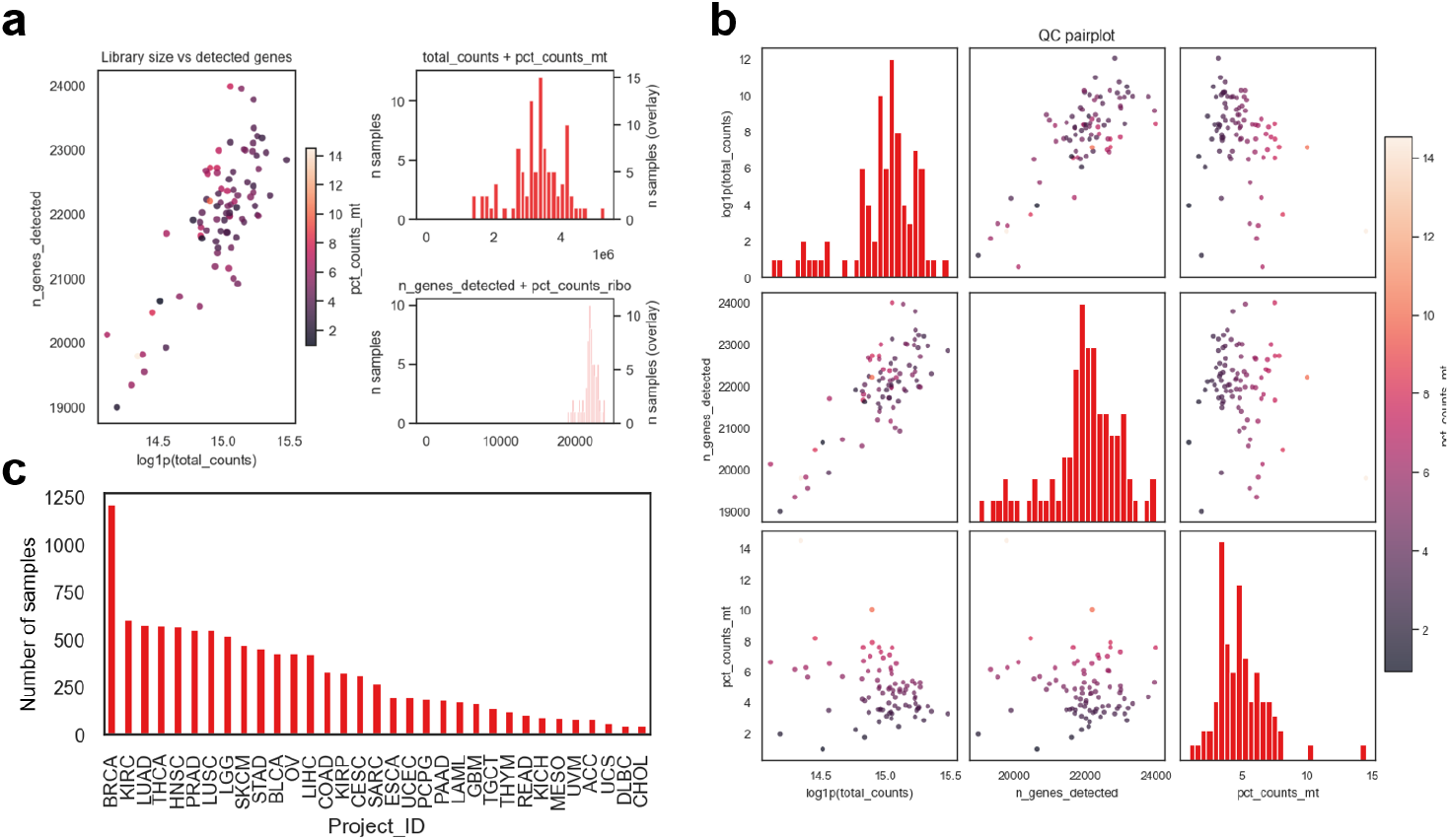
Quality control and sample composition of TCGA bulk RNA-seq data processed with BULLKpy. (**a**) Data were structured into an AnnData object using AnnData and pandas utilities and additional metadata were combined using bk.add_metadata(). Quality control metrics were computed with bk.pp.qc_metrics() and visualized using bk.pl.qc_metrics(). Left, relationship between library size (log_1p_ total counts) and the number of detected genes per sample, colored by mitochondrial read fraction. Right, marginal distributions of total counts and detected genes with overlays of mitochondrial and ribosomal content, enabling identification of technical outliers. (**b**) Pairwise comparison of quality control features generated with bk.pl.qc_pairplot(). Scatter plots illustrate relationships between library size, gene detection, and mitochondrial content, while diagonal histograms show marginal distributions of each metric, providing a global view of sample-level QC variability. (**c**) Distribution of samples across TCGA tumor types after gene and samples were filtered for QC parameters using bk.pp.filter_genes() and bk.pp.filter_samples().

Principal Component Analysis in the TCGA dataset using highly variable genes (Figure 3a) indicated strong variance ratio in PC1 and PC2 (Figure 3b) and we extended routine pipelines by identifying genes with heavy weights in all PCA spaces (Figure 3c,d). Rapid inspection of PCA1 loadings identified neurodevelopmental genes that were later confirmed by plotting their expression in the PCA bidimensional representation (Figure 3e). Further analysis of the pathways affected by these genes can be later done using pathway analysis and gene associations and correlations described below. In addition to transcript expression levels, PCA scatter plots allow the representation of both categorical (e..g. histological grade; Figure 3f) or numerical (tumor purity; Figure 3f) metadata. We also computed standard UMAP (Uniform Manifold Approximation and Projection for Dimension Reduction; (McInnes *et al*, 2018)) plots, as well as the UMAP_graph version (Figure 3g,h). While standard UMAP offers a clean low-dimensional representation, UMAP_graph explicitly displays the sample–sample neighborhood graph (kNN connectivities) underlying the embedding. This reduces overinterpretation of global distances and highlights transitional connectivity patterns relevant to tumor heterogeneity and lineage plasticity.

**Figure 3.**
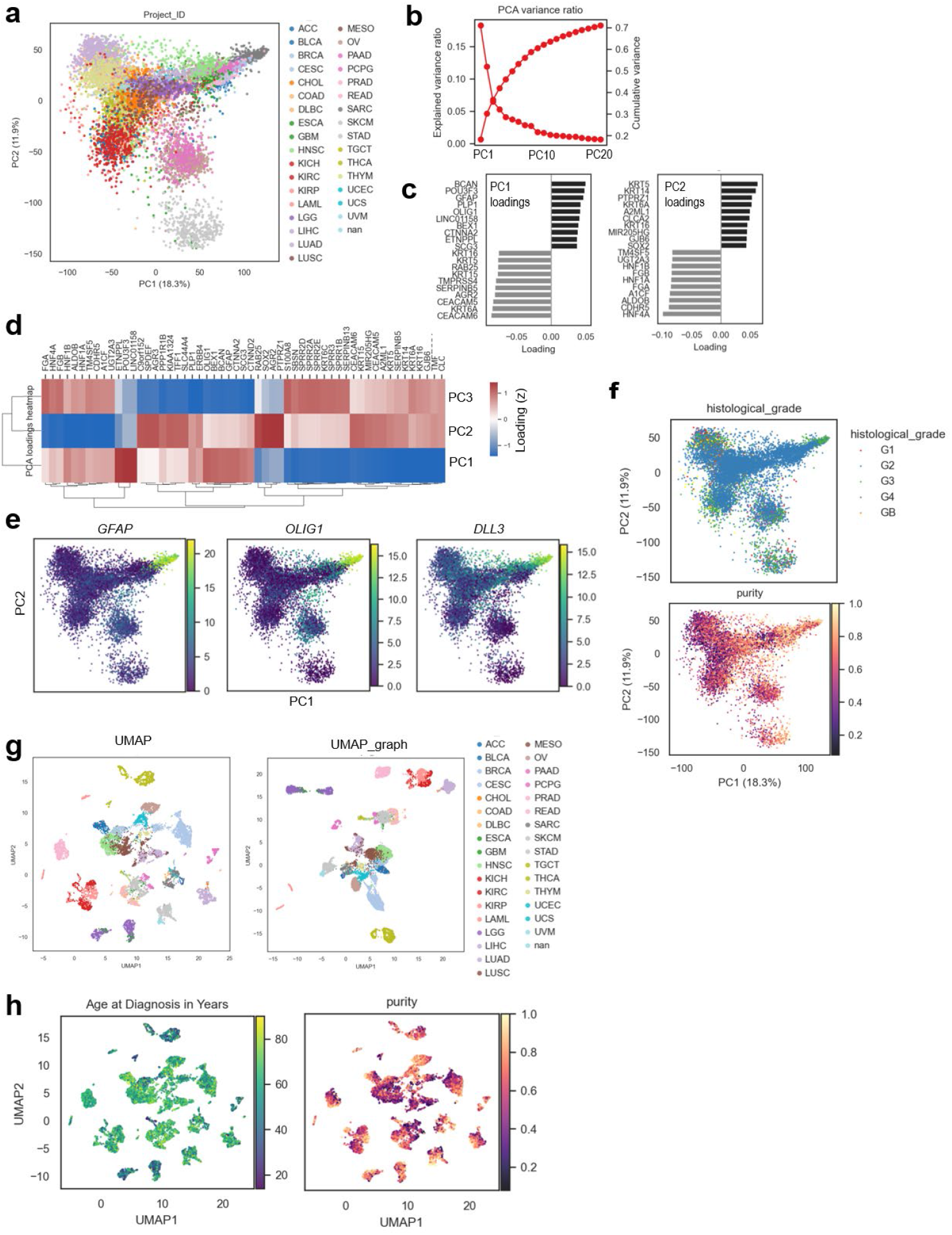
Global transcriptomic structure of TCGA bulk RNA-seq samples revealed by BULLKpy. (**a**) Principal component analysis (PCA) of bulk RNA-seq samples colored by TCGA project, generated using bk.pp.highly_variable_genes() and bk.tl.pca() and visualized with bk.pl.pca_scatter(). The first two principal components capture major sources of transcriptional variation across tumor types. (**b**) Variance explained by principal components, visualized using bk.pl.pca_variance_ratio(). The plot shows both individual and cumulative variance explained, guiding selection of components for downstream analyses. (**c**) Gene loadings for the first two principal components, obtained from bk.tl.pca_loadings() and displayed using bk.pl.pca_loadings_bar(). Top contributing genes highlight biologically interpretable transcriptional programs underlying PCA1 and PCA2. (**d**) Heatmap of standardized gene loadings across the first three principal components, generated with bk.pl.pca_loadings_heatmap(). Hierarchical clustering of genes reveals coordinated contributions to multiple components. (**e**) Expression of selected lineage- and differentiation-associated genes projected onto the PCA space, visualized with bk.pl.pca_scatter(). Gradients along PCA1 and PCA2 illustrate continuous variation in neuroglial and neuroendocrine markers. (**f**) Association of PCA structure with clinical and molecular annotations, shown using bk.pl.pca_scatter() colored by histological grade (top) and tumor purity (bottom). These projections link transcriptional variation to tumor differentiation and cellular composition. (**g**) Uniform Manifold Approximation and Projection (UMAP) embeddings computed from PCA space using bk.tl.neighbors() and bk.tl.umap() and visualized with bk.pl.umap(). Right, bk.tl.umap_graph() overlays the k-nearest-neighbor connectivity graph, emphasizing local neighborhood structure and transitional relationships between tumor groups. (**h**) UMAP projections colored by continuous clinical variables, including age at diagnosis and tumor purity, generated using bk.pl.umap(). These visualizations demonstrate the integration of transcriptomic embeddings with patient-level clinical metadata.

When analyzing new data, clustering of samples by molecular features is a critical discovery step, as it may reveal molecular connection between samples, common biomarkers, etc. We introduced ARI (Adjusted Rand Index; (Hubert & Arabie, 1985)) as a useful statistical approach for a chance-corrected, quantitative measure of agreement between two clusterings (Figure 4a). We also implemented both graph-based and centroid-based clustering approaches. Specifically, we used the Leiden community-detection algorithm (Traag *et al*, 2019) or the k-nearest-neighbor (kNN; (MacQueen, 1967)) graph constructed from PCA-reduced expression profiles, enabling flexible detection of clusters at multiple resolutions (Figure 4b,c). To quantify associations between unsupervised cluster assignments and categorical clinical or molecular annotations, we constructed contingency tables using the pandas.crosstab function, and the strength of association between categorical variables was assessed using Cramér’s V, a normalized effect-size measure derived from the χ^2^ statistic that is robust to differences in sample size and category number (Figure 4d,e).

**Figure 4.**
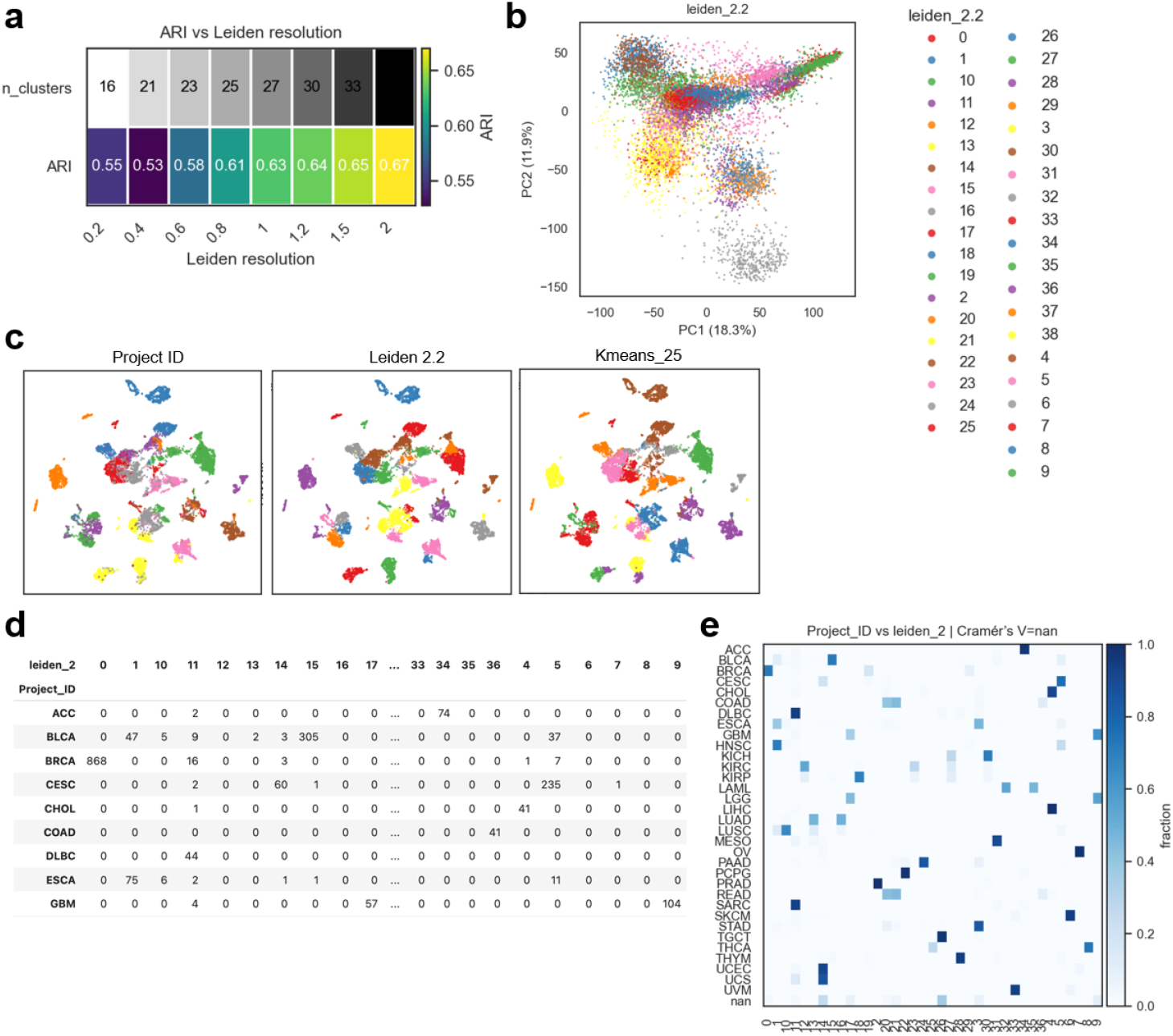
Evaluation and comparison of clustering strategies using BULLKpy. Clustering robustness, concordance with tumor annotations, and correspondence between graph-based and centroid-based approaches were systematically evaluated using quantitative and visual metrics. (**a**) Stability of Leiden clustering across resolutions assessed using the Adjusted Rand Index (ARI), computed with bk.tl.leiden_resolution_scan() and bk.tl.cluster_metrics() and visualized using bk.pl.ari_resolution_heatmap(). ARI values quantify agreement between clustering solutions and reference annotations while accounting for chance, with the corresponding number of clusters indicated for each resolution. (**b**) PCA projection of samples colored by Leiden clusters at resolution 2.2, generated using bk.tl.cluster(method=“leiden”) and visualized with bk.pl.pca_scatter(). This representation illustrates the transcriptional coherence and spatial separation of graph-based clusters in low-dimensional space. (**c**) UMAP embeddings colored by tumor project, Leiden clustering (resolution 2.2), and k-means clustering (k = 25), computed using bk.tl.umap(), bk.tl.cluster(method=“leiden”), and bk.tl.cluster(method=“kmeans”), and visualized with bk.pl.umap(). Side-by-side comparison highlights similarities and differences between clustering approaches and their relationship to tumor origin. (**d**) Contingency table summarizing the overlap between TCGA project annotations and Leiden clusters, generated using pandas.crosstab(). Counts indicate the number of samples shared between each project and cluster. (**e**) Heatmap of cluster–annotation associations showing the fraction of samples per project assigned to each Leiden cluster, computer and visualized with bk.tl.categorical_confusion() and bk.pl.categorical_confusion(). Association strength between cluster labels and tumor type was quantified using Cramér’s V, providing a normalized measure of categorical correlation.

Most of the data generated during these preprocessing and initial evaluation steps can be stored in a stable manner in the AnnData by defining new AnnData observarion (samples) columns or unstructured data (e.g. genes per sample, PCA/UMAP position of each samples, or classification into specific leiden groups).

### Data exploration and gene/metadata correlations and associations

BULLKpy implements many of the most-typically-used quantifications and plots for data exploration including violin plots (Figure 5a,b) or dotplots (Figure 5c) that are able to summarize data both visually and/or using appropriate statistical comparisons. Dot-plot summaries of neuroendocrine and proliferation-related genes across tumor types revealed coherent co-expression patterns, with specific cancers showing strong enrichment of neuroendocrine markers alongside suppression of proliferative signals (Figure 5c). We also implemented the generation of new scores (Figure 5d-f), stored as new observations columns, based on the expression of multiple genes belonging to the same biological process, by implementing algorithms previously used in single-cell pipelines (Hao *et al*, 2024; Wolf *et al*., 2018). A further utilities of these scores is not only generating numeric values for each score per sample but also classifying samples in different groups, as represented by the cell cycle phase classification (Figure 5g) used here as a surrogate of poor or highly-proliferating tumors.

**Figure 5.**
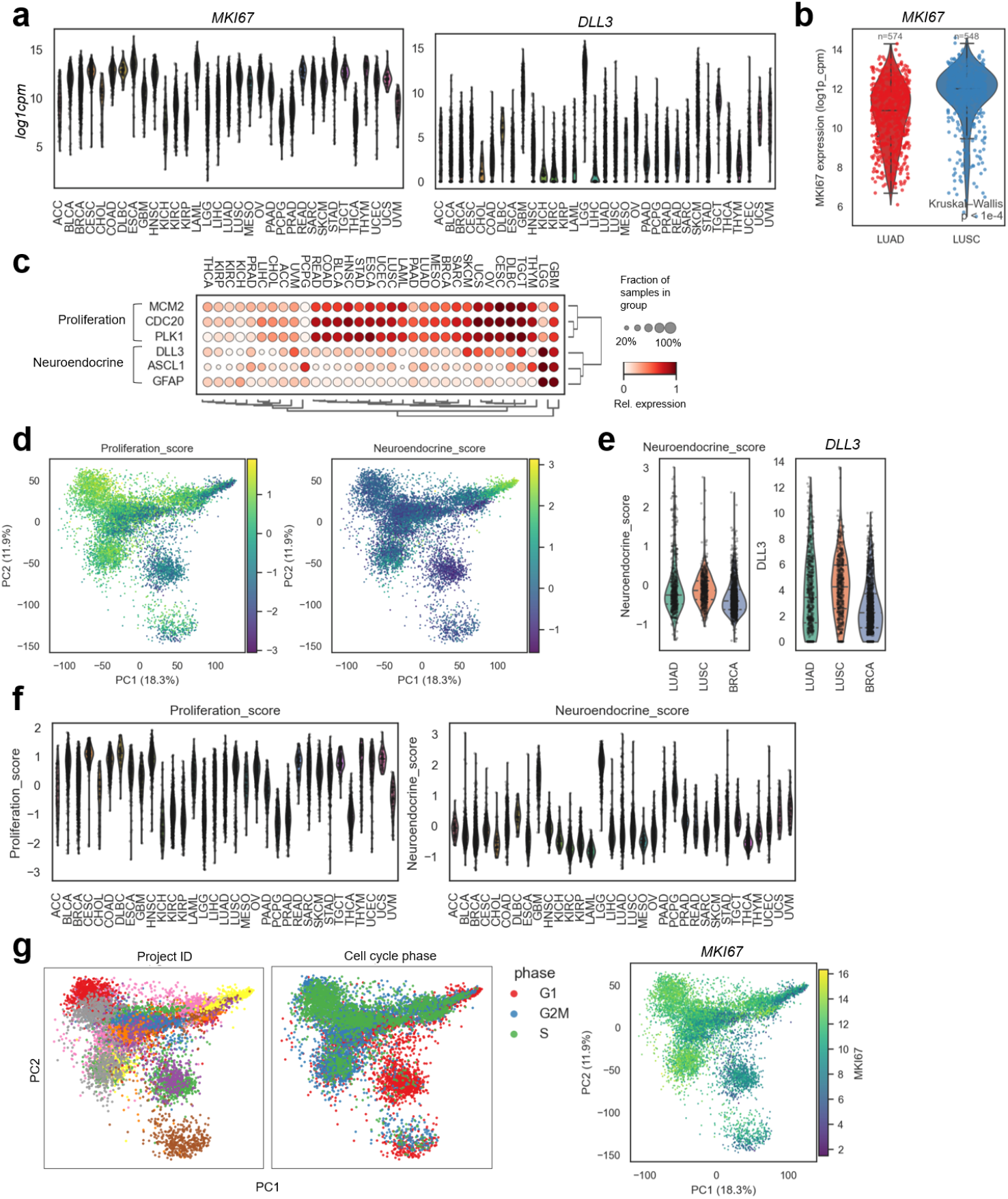
Transcriptional programs of proliferation and neuroendocrine differentiation across tumor types. (**a**) Violin plots showing expression of the proliferation marker *MKI67* and the neuroendocrine marker *DLL3* across TCGA tumor types, visualized using bk.pl.violin().(**b**) Comparison of *MKI67* expression between LUAD and LUSC samples, including statistical analysis visualized with bk.pl.gene_association(). Sample counts are indicated above each group, and statistical significance was assessed using a Kruskal–Wallis test. (**c**) Dot plot summarizing relative expression of selected proliferation and neurodevelopmental genes across tumor types. Dot size indicates the fraction of samples expressing each gene, and color encodes scaled expression, generated with bk.pl.dotplot(). (**d**) PCA projections colored by proliferation and neuroendocrine signature scores, respectively. Scores were computed using bk.tl.score_genes() and visualized on the PCA space with bk.pl.pca_scatter(), highlighting opposing transcriptional gradients along PC1. (**e**) Violin plots of neuroendocrine signature scores and *DLL3* expression across selected tumor types (LUAD, LUSC, BRCA), generated using bk.pl.violin(), illustrating tumor-specific enrichment of neuroendocrine features. (**f**) Distribution of proliferation and neuroendocrine scores across all tumor types, visualized using bk.pl.violin(), summarizing inter-tumor heterogeneity of these functional programs. (**g**) PCA projections colored by tumor project, inferred cell cycle phase (G1, S, G2M), and *MKI67* expression. Cell cycle phases were assigned using bk.tl.score_genes_cell_cycle() and visualized with bk.pl.pca_scatter(), demonstrating the strong association between proliferative state, *MKI67* expression, and PC1.

Biomedical samples are typically accompanied by a large set of metadata (hundreds of sample- or patient-associated metadata in the case of TCGA). We made an effort in BULLKpy to implement additional functions to study gene-gene, gene-metadata and metadata-metadata correlations and associations (numeric and categorical metadata). An example of the analysis of genes correlating with the neuronal marker *DLL3* is shown in Figure 6a, and this analysis can be easily extended for correlating genes with numeric metadata (e.g. *DLL3* versus immune infiltration) or two numeric observations (e.g. immune infiltration and survival). The positive or negative association between genes and specific numeric and categorical metadata is also represented in Figure 6b. Examples of correlation between specific signatures generated above (proliferation and neuroendocrine signatures) and clinical or other numerical metadata are shown in Figure 6c and the corresponding statistical values (R, pval, qval) are also provided during the calculation. Scatter plots stratified by tumor type further illustrated that global correlations can mask subtype-specific behaviors (Figure 6d–f). In lung cancers, for instance, the relationship between neuroendocrine scores and proliferation differed markedly between adenocarcinoma and squamous carcinoma, highlighting the importance of stratified analyses by known groups or newly resolved clusters (Figure 4) when interpreting bulk transcriptomic data.

**Figure 6.**
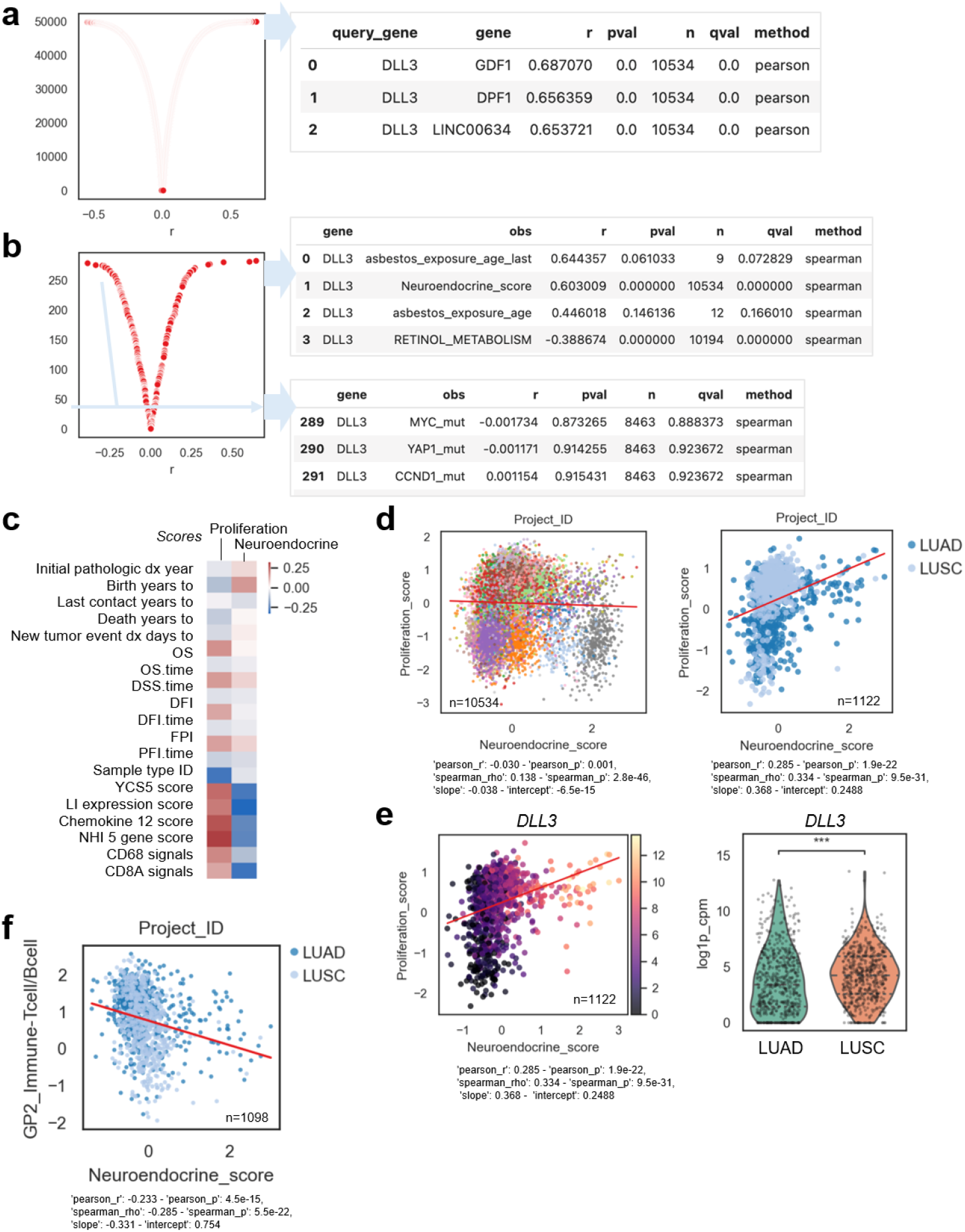
Integrated correlation analysis linking gene expression, transcriptional programs, and clinical variables across TCGA tumors. (**a**) Global gene–gene correlation analysis for the neuroendocrine marker *DLL3*. The left panel shows the distribution of correlation coefficients (r) across all genes, while the table on the right lists the top positively correlated genes ranked by Pearson correlation. Correlations were computed using bk.tl.gene_gene_correlations(). (**b**) Gene–phenotype and gene–mutation association analysis for *DLL3*. Correlations between *DLL3* expression and selected clinical variables, transcriptional scores, and mutation status are shown using Spearman correlation. Results were generated with bk.tl.top_gene_obs_correlations(), enabling simultaneous testing across heterogeneous metadata types. (**c**) Heatmap summarizing correlations between transcriptional scores (Proliferation and Neuroendocrine programs) and key clinical and immune-related variables, including survival endpoints and immune infiltration signatures. Correlation coefficients were calculated using bk.tl.obs_obs_corr_matrix() and visualized with bk.pl.plot_corr_heatmap(). (**d**) Scatter plots showing the relationship between Neuroendocrine and Proliferation scores across all samples (left) and within lung adenocarcinoma and squamous cell carcinoma (right). Linear regression lines and Pearson/Spearman statistics are overlaid. Analyses were performed and visualized using bk.pl.corrplot_obs(). (**e**) Association between *DLL3* expression and transcriptional programs. Left: scatter plot of *DLL3* expression versus Proliferation score with linear regression and correlation statistics using bk.pl.corrplot_obs(). Right: violin plots comparing *DLL3* expression between LUAD and LUSC samples (using bk.pl.gene_association()). (**f**) Relationship between Neuroendocrine score and immune-related metaprogram activity (GP2_Immune–Tcell/Bcell) within lung tumors, highlighting inverse associations between neuroendocrine differentiation and immune infiltration. Correlations were computed and visualized using bk.pl.corrplot_obs().

The analysis of DNA mutations is still leading the efforts in understanding cancer biology and opportunities for treatment in cancer patients. We also implemented oncoprints in which specific mutations are mapped to sample groups (Figure 7a) or expression profiles (Figure 7b). The presence or absence of mutations in samples is encoded as additional obs metadata, and oncoprints provide a compact representation of mutation frequencies, distribution per any selected group or classification of samples, and co-occurrence with other mutations (Figure 7a).

**Figure 7.**
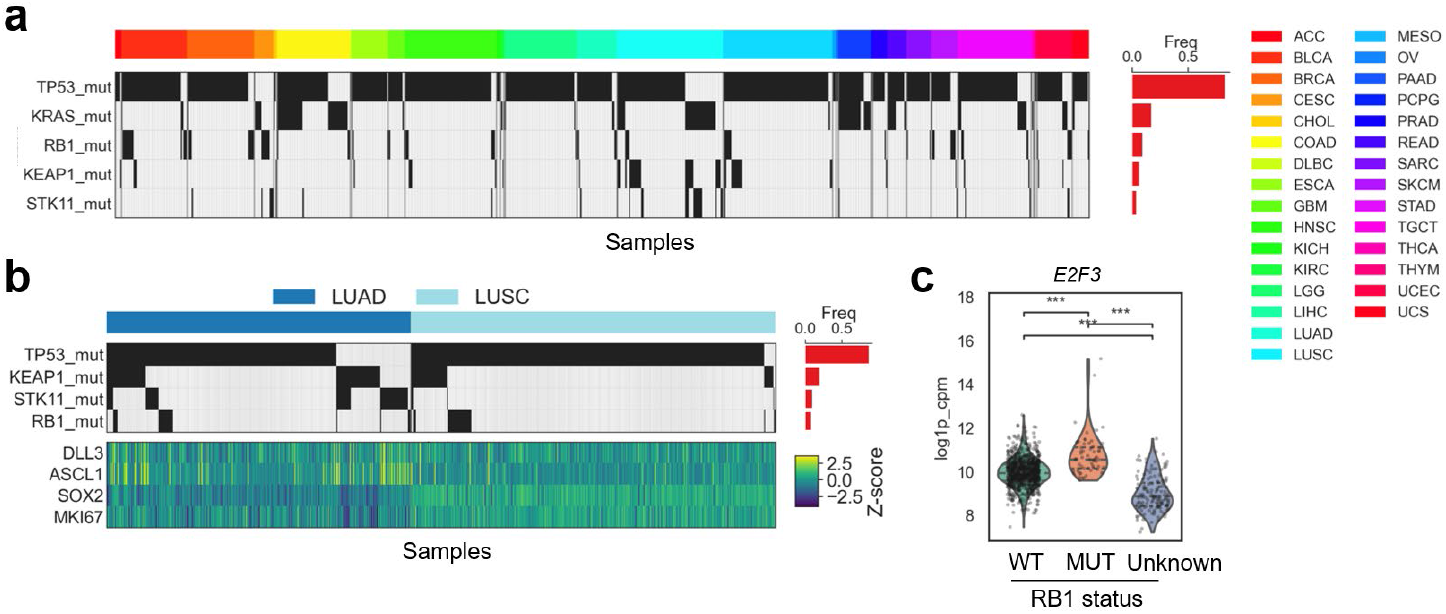
Integrated oncoprint and expression analysis links genomic alterations to transcriptional programs. (**a**) Pan-cancer oncoprint summarizing the mutation status of key cancer driver genes (*TP53, KRAS, RB1, KEAP1* and *STK11*) across TCGA samples, grouped by tumor type. The top color bar indicates tumor origin, and right-side bars show mutation frequencies per gene. (**b**) Tumor-specific oncoprint focusing on lung adenocarcinoma (LUAD) and lung squamous cell carcinoma (LUSC). The upper panel shows mutation status of major drivers, while the lower panel displays z-scored expression of selected genes associated with proliferation (*MKI67*) and neuroendocrine differentiation (*DLL3, ASCL1, SOX2*). In (a,b), mutation and expression matrices were integrated using bk.pl.oncoprint() with combined binary and continuous tracks. (**c**) Differential expression of *E2F3* stratified by *RB1* mutation status (wild-type, mutant, or unknown). Violin plots show log1p-transformed expression values, with statistical significance assessed using a Kruskal–Wallis test. This analysis was performed with bk.pl.gene_association().

In lung cancers, alterations in *TP53, RB1, KEAP1*, and *STK11* displayed characteristic distributions that aligned with neuroendocrine differentiation and proliferative activity (Figure 7b). Simultaneous visualization of mutation status and gene expression highlighted cases where transcriptional programs diverged from canonical genomic expectations, suggesting alternative regulatory mechanisms and effects in downstream targets as represented by the strong association between RB1 mutations and overexpression of E2F transcription factors (Figure 7c).

### Transcriptional programs and their functional and clinical implications

Building on the low-dimensional structure and integrative exploratory analyses described above, we implemented various differential expression and pathway-level analyses tools to translate global transcriptional patterns into mechanistic and clinically interpretable insights. We first performed differential expression analyses across tumor entities or new groups detected after clustering, to identify genes (possible biomarkers) driving the major axes of transcriptional variation per group (Figure 8a,b). We next focused on the differences between the two types of lung cancer represented in TCGA, lung adenocarcinoma (LUAD) and lung squamous cell carcinoma (LUSC), and performed differential expression analysis between these two groups. Data are shown as volcano plots (fold change versus statistical significance; Figure 8c), ranked genes (Figure 8d) or MA (fold change versus mean normalized expression; Figure 8e) plots.

**Figure 8.**
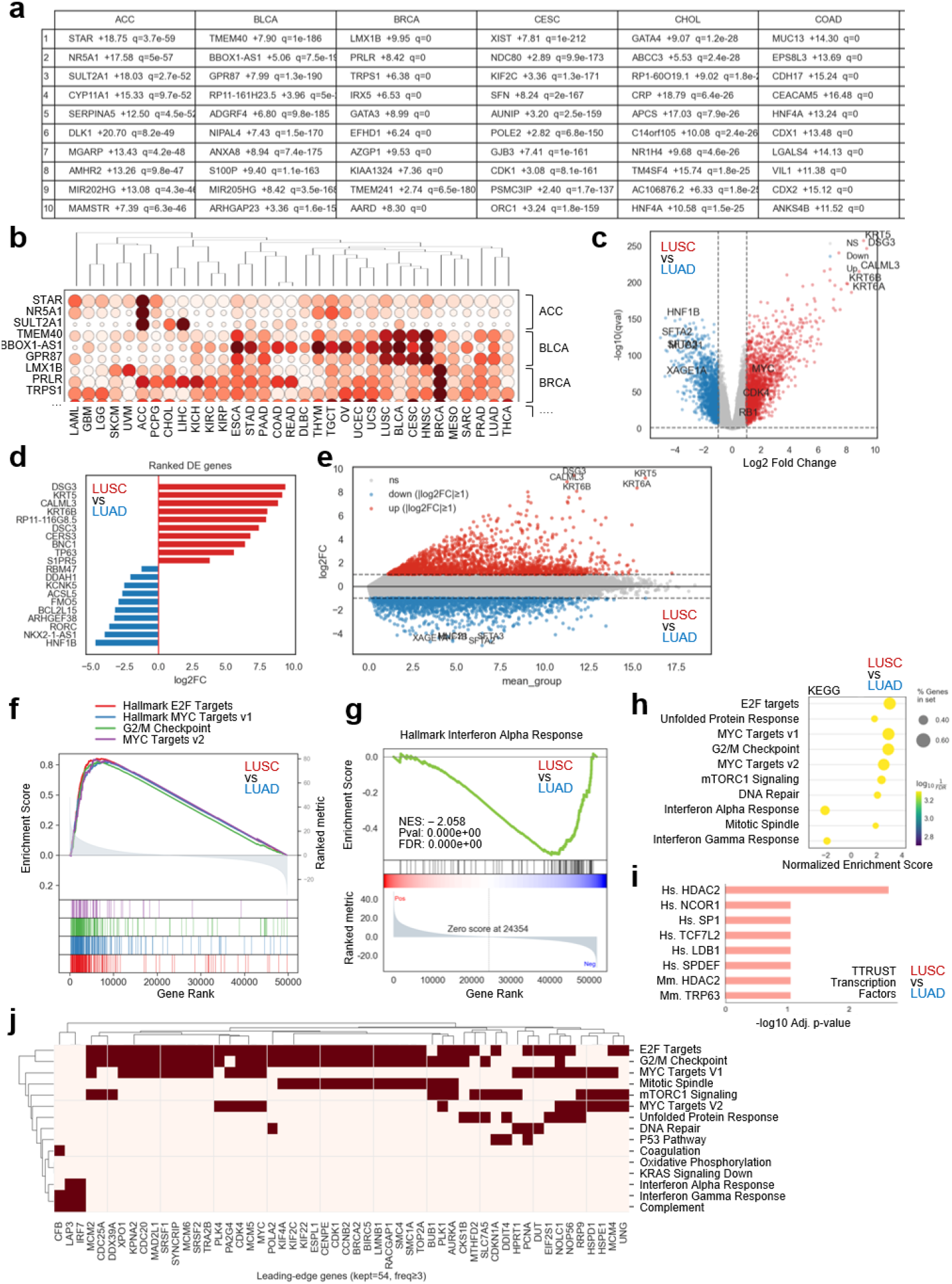
Differential expression and pathway analysis reveal proliferative and neuroendocrine programs distinguishing lung cancer subtypes. (**a**) Summary of top differentially expressed genes across multiple TCGA tumor types. For each cancer type, the top ten genes ranked by effect size and false discovery rate are shown. Differential expression was computed using bk.tl.rank_genes_groups() and summarized with bk.pl.rank_genes_groups(). (**b**) Dot heatmap showing expression patterns of selected differentially expressed genes across tumor types, with hierarchical clustering applied to both genes and cancer types. Dot size represents the fraction of samples expressing each gene, and color intensity reflects relative expression. This visualization was generated using bk.pl.rank_genes_groups_dotplot(). (**c**) Volcano plot comparing lung squamous cell carcinoma (LUSC) versus lung adenocarcinoma (LUAD). Genes are colored by differential expression status (upregulated, downregulated, or non-significant), highlighting key transcriptional drivers such as *MYC, CDK4*, and *RB1*. Differential expression was computed with bk.tl.de() and visualized using bk.pl.volcano(). (**d**) Ranked bar plot of the most differentially expressed genes between LUSC and LUAD, showing direction and magnitude of log2 fold change. This panel was generated using bk.pl.rankplot() based on results from bk.tl.de(). (**e**) MA plot [bk.pl.map()] illustrating the relationship between mean expression and log2 fold change for LUSC versus LUAD, highlighting significantly deregulated genes. (**f**) Gene set enrichment analysis (GSEA) enrichment curves for hallmark pathways associated with proliferation and cell cycle control (E2F targets, MYC targets, G2/M checkpoint). (**g**) GSEA enrichment plot for the interferon alpha response pathway, showing negative enrichment in LUSC compared with LUAD. Enrichment score, normalized enrichment score (NES), and false discovery rate are indicated. (**h**) Summary bubble plot of enriched KEGG pathways, where dot size represents the number of leading-edge genes and color reflects normalized enrichment score. (**i**) Transcription factor enrichment analysis of leading-edge genes using TRRUST, highlighting regulators such as HDAC2, NCOR1, and SP1. Plots f-I were generated by connecting the GSEApy pipeline with the BULLKpy AnnData object and the results from the bk.tl.de() analysis. (**j**) Heatmap of leading-edge genes shared across multiple enriched pathways, illustrating convergence on cell cycle, MYC, DNA repair, and stress response programs. Analysis performed by bk.pl.leading_edge_jaccard_heatmap() and bk.pl.leading_edge_overlap_matrix().

To explore the integration of BULLKpy with external python tools, we applied gene set enrichment analysis to ranked differential expression results using the GSEApy wrap (Fang *et al*, 2023) connected to MDsigDB (Liberzon *et al*, 2015) and enrichr (Chen *et al*, 2013; Xie *et al*, 2021) signatures and data sets. This revealed a coherent set of pathways consistently associated with the observed transcriptional differences, including upregulation of cell proliferation signatures and downregulation of immune transcriptional programs in LUSC when compared to LUAD (Figure 8f,g,h). Specific transcription regulators of the HDAC2, NCOR1, TCF7 and TP63 families were significantly linked to genes upregulated in LUSC (Figure 8i).

We also implemented GSEA leading-edge analyses to identify the core subset of genes recurrently contributing to enrichment across multiple pathways. This scarcely used but critical analysis provides detailed information at the gene level that typically hides behind pathway names, explaining the co-appearance of different pathways sharing a few genes or informing of the specific regulators, within a general pathway, responsible for the differences observed. A quick analysis of differences between LUSC and LUAD suggested three basic clusters of pathways represented by immune genes, E2F-MYC_v1 targets involved in the cell cycle, and MYC_v2 targets related to mTOR, translation and protein processing (Figure 8j).

We also computed the representation in TCGA samples of cellular metaprograms previously reported in a large single-cell curated cancer atlas (Gavish *et al*, 2023). Although these metaprograms were described to compute intratumor heterogeneity, correlation analyses between metaprogram heterogeneity, dispersion, and previously computed scores (proliferation, neuroendocrine, etc.) revealed distinct patterns across tumor types (Figure 9a-c). We first quantified multiple complementary properties of metaprogram organization (Figure 9a), and average metaprogram activity patterns across tumor types (Figure 9b). We then we assessed within–tumor type variability by measuring how strongly metaprogram activity fluctuates among samples of the same cancer type (Figure 9c). Tumor types displayed distinct dispersion profiles, with some cancers showing highly stable program activity across patients, while others exhibited pronounced variability, consistent with increased transcriptional heterogeneity.

**Figure 9.**
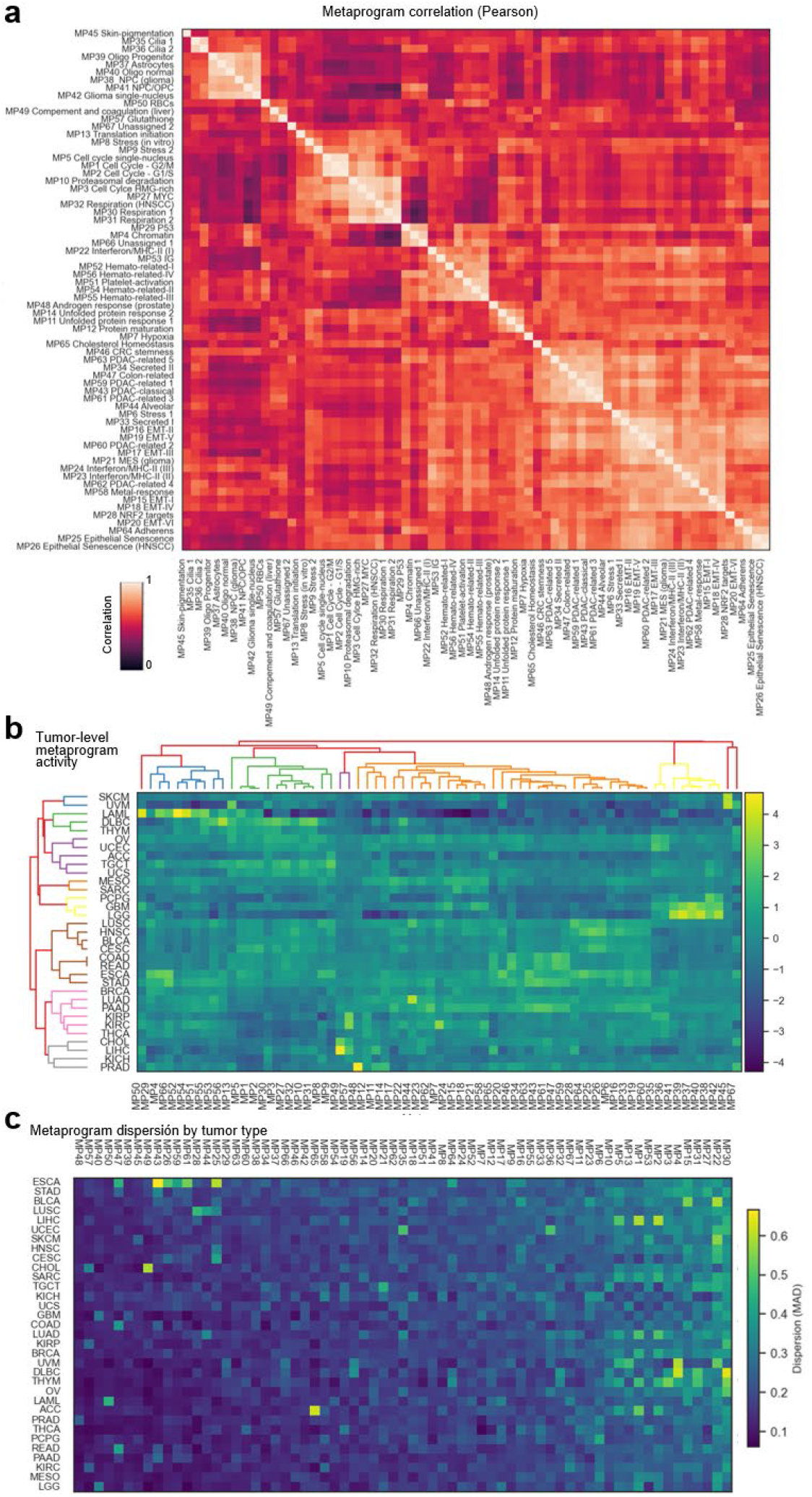
Metaprogram structure, dispersion, and dominance across tumor types. (**a**) Pairwise correlation matrix of metaprogram activities across all samples, computed in bk.tl.score_metaprograms(). Metaprograms cluster [bk.pl.metaprogram_corr()] into coherent biological blocks reflecting shared functional programs such as proliferation, stress response, or immune signaling. (**b**) Heatmap of metaprogram dispersion across tumor types, quantified as median absolute deviation (MAD) of metaprogram weights. This representation by bk.pl.metaprogram_dispersion_heatrmap() highlights tumor-specific variability and plasticity of transcriptional programs. (**c**) Z-scored metaprogram activity profiles across tumor types, with hierarchical clustering applied to both metaprograms and cancers. This analysis reveals tumor-type–specific metaprogram signatures and shared transcriptional states across histologies. Z-scoring and aggregation were performed using bk.pl.metaprogram_heatmap().

We next determined which metaprogram was most prominent in each individual sample and summarized these assignments at the tumor-type level (Figure 10a). This analysis revealed that certain cancers are largely driven by a small number of recurrent programs, whereas others show a more diverse mixture of dominant states across patients. We also quantified the strength of transcriptional dominance by examining how strongly the leading metaprogram outweighs alternative programs within each sample (Figure 10b). To further quantify transcriptional complexity at the program level, we measured per-sample metaprogram heterogeneity and dispersion across tumor types (Figure 10c). Most cancers exhibited high overall metaprogram entropy, indicating that multiple transcriptional programs contribute to individual tumor profiles. Measures of metaprogram dispersion revealed substantial variability in how strongly programs fluctuated across samples within the same tumor type, highlighting differences in intratumoral transcriptional plasticity. Proliferation scores showed a non-linear relationship with metaprogram entropy, with highly proliferative tumors often occupying regions of reduced transcriptional diversity (Figure 10d).

**Figure 10.**
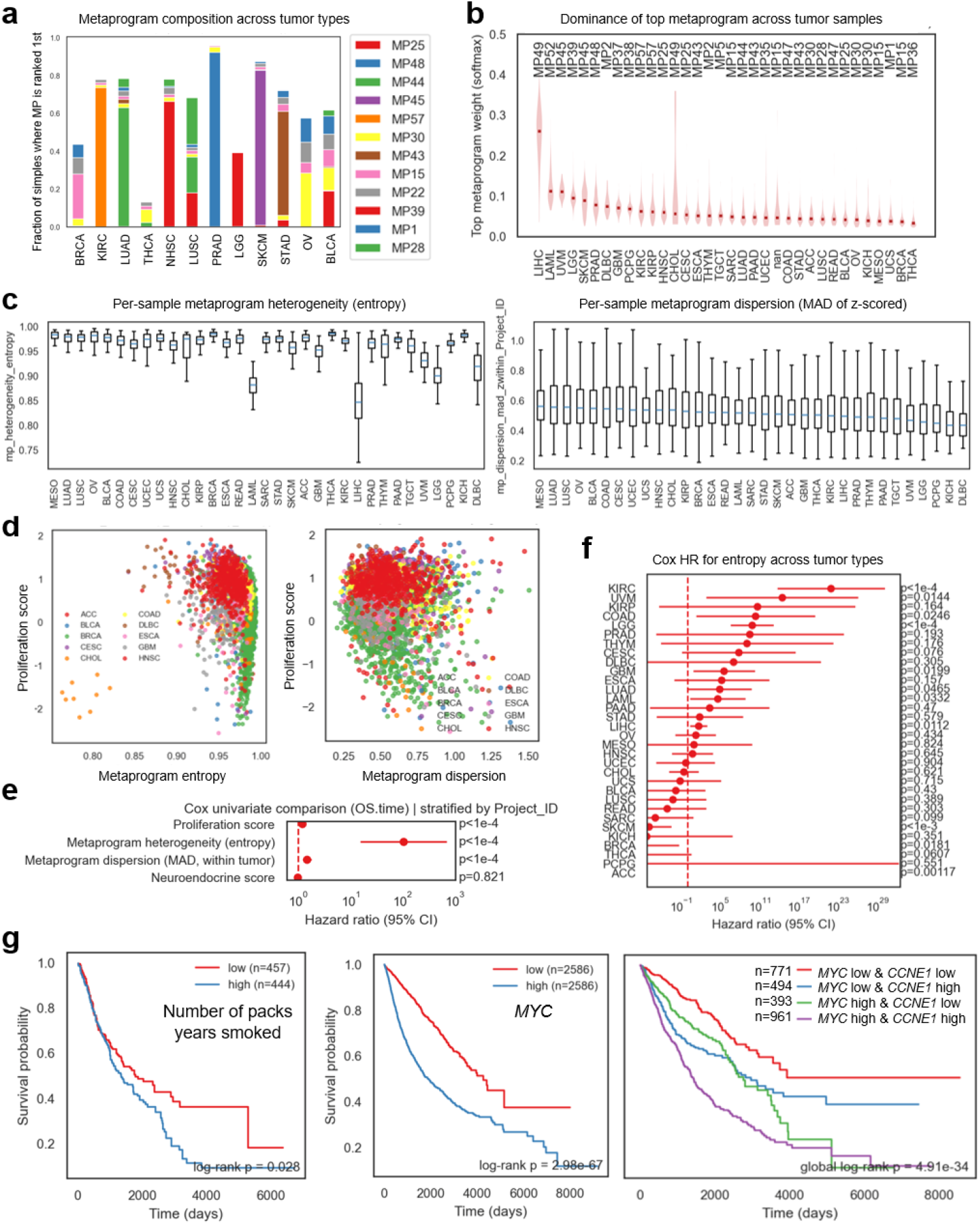
Metaprogram heterogeneity, dispersion, and clinical relevance across tumor types. (**a**) Stacked bar plot showing the composition of rank-1 (dominant) metaprograms across tumor types. Each bar represents the fraction of samples within a tumor type for which a given metaprogram is the top-weighted program, highlighting dominant transcriptional states. Rank-1 assignment was computed using bk.tl.metaprogram_topk_contribution() and visualized with bk.pl.metaprogram_rank1_composition_stackedbar(). (**b**) Violin plot illustrating the dominance of the top-ranked metaprogram across tumor types, quantified as the softmax-normalized weight of the highest metaprogram per sample. Median dominance values are indicated, and metaprogram labels are shown above each violin. This analysis was performed using bk.tl.metaprogram_topk_contribution() and visualized with bk.pl.metaprogram_dominance_ridgeplot_like(). (**c**) Distribution of per-sample metaprogram heterogeneity (left; entropy of metaprogram weights) and per-sample metaprogram dispersion (right; median absolute deviation of z-scored metaprogram activities within tumor types). These metrics capture complementary aspects of transcriptional diversity: heterogeneity reflects the balance of programs within individual tumors, whereas dispersion reflects variability of program activity across samples of the same cancer type. Metrics were computed using bk.tl.metaprogram_metrics_summary() and summarized by tumor type. (**d**) Scatter plots showing the relationship between proliferation program activity and metaprogram heterogeneity (left) or dispersion (right) across individual samples, colored by tumor type. Plots were derived using bk.tl.metaprogram_ne_scatter(). (**e**) Forest plot of univariate Cox proportional hazards models for overall survival (OS), stratified by tumor type, comparing proliferation score, metaprogram heterogeneity (entropy), metaprogram dispersion (within-tumor MAD), and neuroendocrine score. Hazard ratios and confidence intervals were estimated using bk.tl.cox_univariate() with stratification by Project_ID and visualized with bk.pl.cox_forest_from_uns(). (**f**) Tumor-type–specific hazard ratios for metaprogram heterogeneity (entropy), estimated using Cox models stratified by cancer type. This analysis reveals tumor-specific prognostic effects of transcriptional heterogeneity, with significant associations observed in multiple cancer entities. Cox models were computed using bk.tl.cox_by_group() and visualized with bk.pl.cox_forest_by_group(). (**g**) Kaplan–Meier survival analyses illustrating the prognostic impact of smoking (left), *MYC* program activity (middle), and their interaction with *CCNE1* status (right). Patients were stratified into low and high groups using quantile-based binning, and survival differences were assessed using log-rank tests. Survival groups and Kaplan–Meier curves were generated using bk.pl.km_univariate() and bk.pl.km_2×2_interaction().

We also implemented survival analyses using overall survival while accounting for tumor-type–specific baseline risks. In univariate models, both metaprogram heterogeneity and dispersion were significantly associated with patient outcome, with higher transcriptional complexity corresponding to increased hazard (Figure 10e). When the prognostic impact of metaprogram heterogeneity was evaluated across individual tumor types, we observed association between increased entropy and poorer outcome in several cancers, although effect sizes varied (Figure 10f). Finally, we implemented Kaplan–Meier analyses on AnnData either from observations and metadata (e.g. smoking habits) or from gene expression profiles. Kaplan–Meier analyses revealed that both elevated smoking and high *MYC* expression were individually associated with worse survival. In addition, stratification by combined interactions (e.g. *MYC* or/and *CCNE1* expression levels) uncovered additive and synergistic effects, with tumors exhibiting concurrent activation of MYC and cell cycle programs displaying the poorest outcomes (Figure 10g).

## Discussion

Bulk OMICs data remains one of the most widely used technologies in biomedical and cancer research, owing to its robustness, cost-effectiveness, and the vast number of available public datasets accompanied by rich clinical and genomic annotations. Despite the rapid expansion of single-cell technologies, bulk transcriptomic data continue to provide unique advantages, particularly for large cohorts, long-term clinical follow-up, and integrative analyses that combine gene expression with genomic alterations, histopathology, and outcome measures, features exemplified by TCGA pan-cancer resources (Cancer Genome Atlas Research *et al*., 2013; Hoadley *et al*., 2018). However, the analytical ecosystem for bulk OMICs has historically been fragmented, with key functionalities distributed across disparate tools, languages, and data structures. This fragmentation has limited reproducibility, increased technical barriers, and slowed the adoption of integrative workflows.

A central contribution of BULLKpy is the consolidation of widely used and state-of-the-art analytical methods into a single, coherent pipeline built around the AnnData object and the conceptual structure established by the Scanpy ecosystem (Virshup *et al*., 2023; Wolf *et al*., 2018). Scanpy popularized a preprocessing-analysis-visualization workflow for transcriptomics at scale, while AnnData provides a standardized container for expression matrices, metadata, and derived results. By adopting a preprocessing–tools– plotting architecture, BULLKpy enables users to move seamlessly from data integration to exploratory analysis, statistical testing, functional interpretation, and clinical association studies. This design not only reduces the cognitive and technical overhead required to perform complex analyses, but also facilitates transparent and reproducible research practices. By integrating dozens of different analyses in a single notebook, BULLKpy lowers the barrier for researchers transitioning between wet and in silico expertise, and encourages methodological convergence within the python ecosystem.

One of the core interests of BULLKpy is to facilitate a number of statistical tests linking molecular and functional or clinical data. GSEA introduced a framework for interpreting expression changes at the level of predefined gene sets, improving robustness and interpretability when individual-gene signals are subtle or heterogeneous (Subramanian *et al*, 2005). Complementary approaches such as GSVA compute sample-wise pathway activity scores, enabling continuous phenotyping and association testing across cohorts (Hanzelmann *et al*, 2013). Functional interpretation in BULLKpy is strengthened by explicit support for leading-edge prioritization, a test introduced with GSEA that, despite its functional utility, has been poorly explored in functional analysis of OMICs data. BULLKpy also extends bulk RNA-seq analysis toward higher-order organization of transcriptional states via tumor cell metaprogram analysis (Gavish *et al*., 2023). Conceptually, metaprograms relate to module-based frameworks widely used in bulk expression studies, such as co-expression network approaches (e. g., WGCNA), which identify correlated gene modules and relate them to external traits (Langfelder & Horvath, 2008). By decomposing expression matrices into recurrent gene programs and quantifying their activity, dispersion, and heterogeneity across samples, metaprograms provide a compact representation of complex tumor states. Importantly, quantifying not only program activity but also program organization (e.g., dominance, entropy-like measures) creates a bridge between bulk transcriptomics and single-cell-inspired views of plasticity and state mixtures. When combined with survival and clinical metadata, these summaries enable interpretable associations between transcriptional architecture and patient outcome, supporting both hypothesis generation and translational stratification.

A further distinguishing feature of BULLKpy is native support for oncoprint-style visualizations that integrate somatic mutation data with transcriptional and clinical information. Oncoprints are widely used in cancer genomics to summarize alteration patterns across samples and genes, popularized through portals such as cBioPortal and implemented in visualization frameworks including ComplexHeatmap (Cerami *et al*, 2012). Yet, oncoprints are often deployed separately from transcriptomic workflows, limiting their use in integrated molecular interpretation. By embedding oncoprints within an expression-centered pipeline, BULLKpy enables direct comparison between genomic alterations and expression-derived phenotypes, including transcriptional programs, novel sample clusters or groups, or further clinical metadata. This integrated perspective is critical for understanding how diverse genomic events converge on shared transcriptional outputs that shape tumor behavior and patient outcomes.

Beyond immediate analytical capabilities, BULLKpy hopes to contribute to a broader vision for computational biology. As machine learning and AI approaches become increasingly central, standardized data structures, interoperable pipelines, and transparent workflows will be essential for reproducibility and tool composability. The scverse initiative was created to address exactly these challenges in the Python ecosystem, emphasizing shared data formats and interoperable APIs (Virshup *et al*., 2023). By positioning bulk OMIC analysis within the same design space as Scanpy/AnnData, BULLKpy aims to help standardize and democratize Python-based transcriptomic workflows, enabling researchers to move more easily between bulk, single-cell, and future multi-modal analyses in a shared ecosystem.

## Methods

### Software architecture and implementation

All analyses were performed using BULLKpy (v.0.0.1), a Python package developed for comprehensive bulk OMICs data analysis with a particular initial focus on cancer transcriptomics. BULLKpy is implemented in Python (v.3.10.10) and distributed as an open-source package via GitHub (https://github.com/malumbres/BULLKpy), with comprehensive documentation available through Read the Docs (https://bullkpy.readthedocs.io/en/latest/index.html). The package is designed to integrate seamlessly into standard Python scientific environments, relying on widely adopted libraries including NumPy (v. 1.23.5), Pandas (v. 2.3.3), Matplotlib (v. 3.8.0), and Seaborn (v. 0.11.2). This choice of dependencies ensures long-term maintainability and compatibility with existing bioinformatics workflows.

The source code is organized into modular subpackages (Figure 1) corresponding to preprocessing (bullkpy.pp), analytical tools (bullkpy.tl), and plotting utilities (bullkpy.pl). The core principle of BULLKpy is the AnnData object (annadata v. 0.10.3), a unified and flexible data structure that tightly couples expression matrices with rich sample- and feature-level metadata, as well as multiple layers of derived representations and results, within a single container (Virshup *et al*, 2024; Wolf *et al*., 2018). All functions operate directly on AnnData objects and store intermediate and final results within the object’s structured fields (.obs,.var,.obsm,.uns), promoting transparency and reproducibility. Version-controlled development, unit testing, and continuous documentation updates are used to ensure robustness and extensibility of the framework. Jupyter notebooks (Jupyter Lab v4.0.2) are used for version control and reproducibility.

### Overview of major analytical modules

The preprocessing module (pp) implements bulk RNA-seq–specific normalization and transformation strategies, including counts-per-million scaling, log-transformation, gene filtering, and variance-based feature selection. These steps are designed to be conceptually analogous to single-cell preprocessing while respecting the statistical properties of bulk data. The tools module (tl) contains the core analytical functionality of BULLKpy. This includes dimensionality reduction (PCA, neighborhood graphs, UMAP), unsupervised clustering, differential expression analysis, pathway and gene set scoring, gene–metadata association testing, hazard ratios and survival studies from scikit-learn (v.1.6.1), and the inference of transcriptional metaprograms.

GSEA (Subramanian *et al*., 2005) pipelines are implemented by connecting to the external GSEApy (v. 1.1.0) python wrap (Fang *et al*., 2023), which gives access to and the MSigDB signature databases (Liberzon *et al*., 2015) and enrichr datasets (Chen *et al*., 2013). This interaction shows an example possible connection between BULLKpy and other past or future python pipelines for biomedical data.

Overall survival and progression-free survival were analyzed using Kaplan–Meier estimators, and statistical significance between groups was assessed using log-rank tests. Cox proportional hazards regression models were used to estimate hazard ratios and to evaluate the independent prognostic value of transcriptomic signatures, with optional adjustment for clinical covariates such as age, tumor stage, and molecular subtype.

Metaprograms are inferred from expression or functional score matrices derived from the Curated Cancer Cell Atlas (https://www.weizmann.ac.il/sites/3CA/methods)(Gavish *et al*., 2023), and stored in structured AnnData fields, enabling downstream analyses of dominance, heterogeneity, dispersion, and clinical relevance. Metaprogram activity scores were averaged across samples within each tumor type, followed by z-score normalization across tumor types for each metaprogram. Hierarchical clustering was applied independently to tumor types and metaprograms using correlation-based distances, revealing patterns of shared transcriptional programs across cancers. Dispersion was then quantified for each metaprogram as the median absolute deviation (MAD) across samples within the same tumor type, providing a robust measure of intra-tumoral transcriptional variability. For each sample, the metaprogram with the highest relative weight (rank-1) was identified. Tumor-type–specific compositions were then calculated as the fraction of samples in which each metaprogram was ranked first, providing a discrete summary of dominant transcriptional programs across cancers. For each tumor type, the relative weight of the rank-1 metaprogram was extracted per sample using softmax-normalized metaprogram scores. Distributions of these weights were visualized to quantify the strength and consistency of transcriptional dominance, distinguishing tumors driven by a single prevailing program from those exhibiting more balanced or heterogeneous program usage.

The module also includes cancer-focused methods such as oncoprint-style (Cerami *et al*., 2012) mutation visualization, lineage and differentiation scoring, and survival modeling using Cox proportional hazards and Kaplan–Meier analyses. The plotting module (pl) provides standardized, publication-ready visualizations tightly linked to the underlying analyses. These include heatmaps, ridge and violin plots, stacked bar charts, survival curves, forest plots, and metaprogram-centric visual summaries, all designed on matplotlib and seaborn to operate directly on AnnData-stored results and metadata.

## Availability

The transcriptomic and clinical data analyzed in this study were obtained from publicly available repositories (https://portal.gdc.cancer.gov/; https://xenabrowser.net/). The BULLKpy package is openly available on GitHub at https://github.com/malumbres/BULLKpy, and comprehensive documentation is provided via Read the Docs at https://bullkpy.readthedocs.io/en/latest/index.html. The specific version of BULLKpy used in this study is indicated in the manuscript and in the analysis repository. A complete notebook for this pipeline is available in the GitHub repository (https://github.com/malumbres/BULLKpy/tree/main/notebooks).

## Conflict of Interest

The author declares no competing interests related to this manuscript.

## Acknowledgements

The author wishes to thank first to F.J. Theis, F.A. Wolf and their co-workers (Helmholtz München, Germany) for generating the AnnData and scverse environment. Their work represents a major advance in democratizing single-cell data for the scientific community. The author also thanks all members of the VHIO Cancer Cell Cycle lab for stimulating ideas and discussion. This work was supported by research grants from the Spanish Ministry of Science and Innovation (MICINN), Agencia Estatal de Investigación (AEI)/FEDER (PID2024-161681OB-10 and iDIFFER network RED2024-153635-T), CIBER -Consorcio Centro de Investigación Biomédica en Red-(CIBERONC), Instituto de Salud Carlos III, MICINN, and the SOSCLC-AECC Retos grant from the Asociación Española contra el Cáncer. VHIO would like to acknowledge the Cellex Foundation for providing research facilities and equipment, the CERCA Program from the Generalitat de Catalunya for their support on this research, and funding from MICINN/AEI Centers of Excellence Severo Ochoa (CEX2020-001024-S/AEI/10.13039/501100011033).

## Notes

### Competing Interest Statement

The authors have declared no competing interest.

